# Language model-based B cell receptor sequence embeddings can effectively encode receptor specificity

**DOI:** 10.1101/2023.06.21.545145

**Authors:** Meng Wang, Jonathan Patsenker, Henry Li, Yuval Kluger, Steven H. Kleinstein

**Affiliations:** Program in Computational Biology and Bioinformatics, Yale University, New Haven, CT; Program in Applied Mathematics, Yale University, New Haven, CT; Department of Pathology, Yale School of Medicine, New Haven, CT; Department of Immunobiology, Yale School of Medicine, New Haven, CT, USA

## Abstract

High throughput sequencing of B cell receptors (BCRs) is increasingly applied to study the immense diversity of antibodies. Learning biologically meaningful embeddings of BCR sequences is beneficial for predictive modeling and interpretability. Several embedding methods have been developed for BCRs, but no direct performance benchmarking exists. Moreover, the impact of the input sequence length and paired-chain information on the prediction remains to be explored. We evaluated the performance of multiple embedding models to predict BCR sequence properties and receptor specificity. Despite the differences in model architectures, most embeddings effectively capture BCR sequence properties and specificity. BCR-specific embeddings slightly outperform general protein language models in predicting specificity. In addition, incorporating full-length heavy chains and paired light chain sequences improve the prediction performance of all embeddings. This study provides insights into the properties of BCR embeddings to improve downstream prediction applications for antibody analysis and discovery.

## INTRODUCTION

B cell receptors (BCRs) play a central role in the immune system’s ability to recognize and respond to pathogens. These receptors are expressed on the surface of B cells and directly bind to molecules present on the surface of pathogens, a crucial step in initiating adaptive immune responses. Each B cell expresses a BCR that is practically unique, and this diversity allows the immune system to recognize and respond to any dangerous pathogens. High-throughput sequencing technologies enable large-scale characterization of BCR repertoires, generating massive datasets that can benefit from natural language processing (NLP) methods (Ostrovsky-Berman *et al*., 2021; Leem *et al*., 2022; Ruffolo *et al*., 2021). These NLP methods learn representations (embeddings) for amino acids or groups of amino acids and summarize them across the sequence to create meaningful representations for downstream tasks, such as supervised prediction. Some popular embedding-based models include word2vec (Mikolov *et al*., 2013) and deep transformer models (Vaswani et al., 2017). Immune2vec (Ostrovsky-Berman et al., 2021b) is a word2vec model that learns to represent BCRs as vectors. It does this by breaking down each BCR into smaller units of three amino acids (3-mers), where each unit is embedded into fixed-length sequence representation and then averaged across the sequence to produce a single vector for the given BCR. Recent deep protein transformer models create contextualized embeddings and achieve state-of-the-art performance in downstream prediction tasks, such as secondary structure and protein-protein binding prediction (Lin *et al*., 2022; Elnaggar *et al*., 2020; Filipavicius *et al*., 2020). ESM2 (Lin *et al*., 2022) and ProtT5 (Elnaggar *et al*., 2020) are two examples of transformer models trained on large corpora of protein sequences to create amino acid representations that account for sequence context. To capture the influence of neighboring amino acids on each other, these models generate local embeddings for each amino acid that depend on the whole sequence and compute a global embedding for the entire sequence by averaging the local embeddings. Similar approaches have also been recently applied to immune receptor sequences to train models that predict binding-related properties of immune cell receptors (Leem *et al*., 2022; Wu *et al*., 2021).

The abundance of embedding approaches calls for comparative studies to examine their biological relevance. A critical evaluation objective is how well low-dimensional representations preserve information for downstream prediction tasks (Bengio *et al*., 2014). Previous work observed that neighboring BCRs in the embedding space have similar gene usage and somatic hypermutation frequency (Ostrovsky-Berman *et al*., 2021; Leem *et al*., 2022). However, no quantitative assessment of the representation over prediction tasks exists, and the comparative advantages of each embedding are underexplored. For instance, even though transformer models are highly expressive and can encode complicated context-based relationships, they require more training data to create meaningful and generalizable representations. Models like immune2vec, though less expressive, can be trained on more specific datasets, potentially allowing for a more informative BCR-specific embedding. Here we evaluated multiple embedding methods over prediction tasks, including BCR sequence properties and receptor specificity, to assess how well they preserve biological information.

Previous machine-learning studies on BCR mainly focused on the complementarity-determining region 3 (CDR3) of heavy chain BCR sequences (Ostmeyer *et al*., 2019; Ostrovsky-Berman *et al*., 2021), a determinant of antibody specificity (Xu and Davis, 2000). The recent development of single-cell technologies leads to the increasing availability of paired full-length heavy and light chain BCR sequences, which brings the opportunity to include regions outside CDR and the light chain. However, no studies have examined the effect of incorporating full-length heavy and light chain sequences in receptor specificity prediction tasks.

In this study, we compared the performance of protein language models, including the BCR-specific word2vec model (immune2vec), transformer-based general protein language models (ESM2, ProtT5), and traditional amino acid encoding (physicochemical encoding, amino acid frequency) in predicting BCR sequence properties and receptor specificity. We also examined the effect of incorporating full-length and paired light chain sequences on the prediction performance. We found the BCR-specific immune2vec models perform similarly or slightly outperform general protein language models in receptor specificity prediction tasks. We also found an improvement in specificity prediction performance by incorporating full-length heavy and paired light chain sequences. These observations offer insights into the performance characteristics of embedding methods trained with different types of BCR sequence input and downstream prediction tasks.

## MATERIAL AND METHODS

### Data sources and processing

We collected 1 million single-cell paired heavy and light chain full-length BCR V(D)J sequences from ten datasets (Table 1). In total, 0.87 million unique heavy chain sequences and 0.55 million unique light chain sequences were available from at least 77 donors, excluding CoV-AbDab (Raybould *et al*., 2021) which reported 584 sources/studies. We translated the nucleotide sequences into amino acids using Alakazam 1.0.2 (Gupta *et al*., 2015). The median lengths of the heavy and light chains were 122 and 108 amino acids, respectively.

**Table 1.**
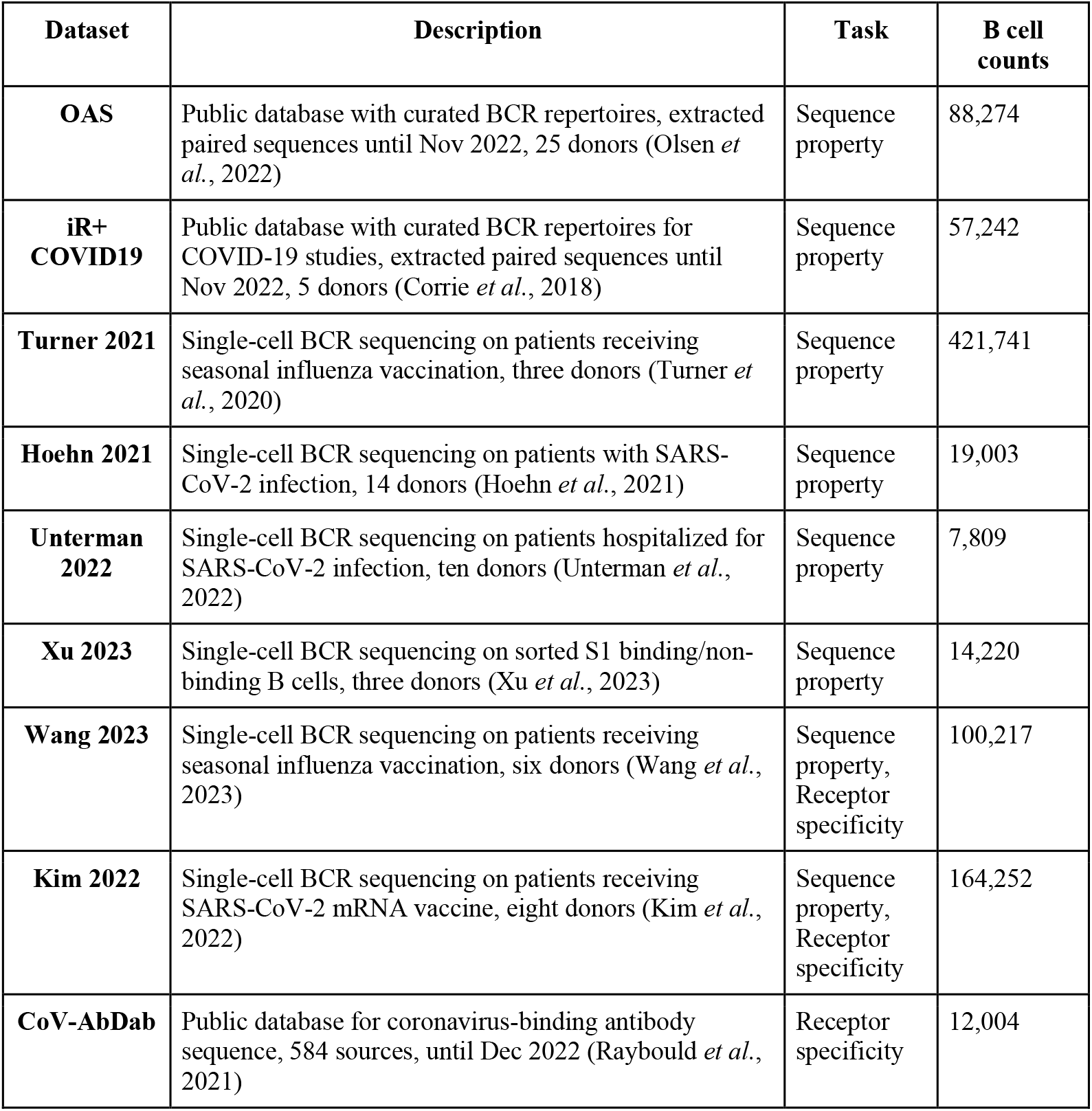
Source and size of the paired BCR heavy and light chain V(D)J sequences datasets used in the study. Each dataset was filtered for B cells with both heavy and light chain sequences.

| Dataset | Description | Task | B cell counts |
| --- | --- | --- | --- |
| **OAS** | Public database with curated BCR repertoires, extracted paired sequences until Nov 2022, 25 donors (Olsen *et al.*, 2022) | Sequence property | 88,274 |
| **iR+ COVID19** | Public database with curated BCR repertoires for COVID-19 studies, extracted paired sequences until Nov 2022, 5 donors (Corrie *et al.*, 2018) | Sequence property | 57,242 |
| **Turner 2021** | Single-cell BCR sequencing on patients receiving seasonal influenza vaccination, three donors (Turner *et al.*, 2020) | Sequence property | 421,741 |
| **Hoehn 2021** | Single-cell BCR sequencing on patients with SARS-CoV-2 infection, 14 donors (Hoehn *et al.*, 2021) | Sequence property | 19,003 |
| **Unterman 2022** | Single-cell BCR sequencing on patients hospitalized for SARS-CoV-2 infection, ten donors (Unterman *et al.*, 2022) | Sequence property | 7,809 |
| **Xu 2023** | Single-cell BCR sequencing on sorted S1 binding/non-binding B cells, three donors (Xu *et al.*, 2023) | Sequence property | 14,220 |
| **Wang 2023** | Single-cell BCR sequencing on patients receiving seasonal influenza vaccination, six donors (Wang *et al.*, 2023) | Sequence property, Receptor specificity | 100,217 |
| **Kim 2022** | Single-cell BCR sequencing on patients receiving SARS-CoV-2 mRNA vaccine, eight donors (Kim *et al.*, 2022) | Sequence property, Receptor specificity | 164,252 |
| **CoV-AbDab** | Public database for coronavirus-binding antibody sequence, 584 sources, until Dec 2022 (Raybould *et al.*, 2021) | Receptor specificity | 12,004 |

For sequence property prediction tasks, we annotated the sequence for V, J gene usage, isotype (IGHM, IGHD, IGHG or IGHA), light chain type (IGK or IGL), somatic hypermutation frequency, and CDR3 length using immcantation 4.3.0 (Gupta *et al*., 2015). We also extracted the SARS-CoV-2 spike protein binding/non-binding labels from datasets for the receptor specificity task. To balance the dataset, we randomly sampled 1,000 sequences from each donor of a pre-COVID-19 dataset (Wang *et al*., 2023) as negatives for specificity prediction.

### Sequence embeddings

#### Immune2vec

We trained four immune2vec models (full-length heavy chain, full-length light chain, CDR3 heavy chain, and CDR3 light chain) with the suggested 100 dimensions (Ostrovsky-Berman *et al*., 2021). We also trained additional immune2vec models with dimensions of 25, 50, 150, 200, 500, and 1,000 to examine the effect of dimensionality. Immune2vec learned the embedding for individual 3-mers that appeared in the sequence corpus and took the average of embeddings for all possible 3-mers along the sequences as sequence-level embedding. For receptor-level embeddings, we concatenated the heavy and light chain sequence-level embeddings.

#### ESM2

We used the pre-trained ESM2 model with 650 million parameters (Lin *et al*., 2022) to generate BCR embeddings. ESM2 outputs embeddings of 1,280 dimensions for individual amino acids within the sequence. We averaged the amino acid embeddings as sequence-level representations.

#### ProtT5

We used the pre-trained ProtT5 (Elnaggar *et al*., 2020) to generate embeddings with a dimension of 1,024 for individual amino acids within the sequence. We averaged the amino acid embeddings as sequence-level representations.

#### Baseline encoding and random control

For sequence property prediction tasks, we used the amino acid frequency (*frequency*) with a dimension of 20 and a collection of physicochemical-based amino acid encoding (*physicochemical*) with 75 latent dimensions from the Python peptides package v0.3.1 (https://pypi.org/project/peptides/) averaged across sequences as baselines. We also evaluated the performance of randomly shuffled embeddings (*shuffled*).

### Prediction tasks

#### V, J gene family (classification)

We framed the gene usage prediction into four separate multiclass classification tasks: given BCR heavy or light chain, predict the V and J gene families used. Classes with smaller than 50 sequences were excluded. The target classes to predict included: Heavy chain V gene: IGHV1 -IGHV7; Heavy chain J gene: IGHJ1 -IGHJ6; Light chain V gene: IGKV1 -IGKV7, IGLV1 -IGLV11; Light chain J gene: IGKJ1 -IGKJ5, IGLJ1 -IGLJ7

#### Heavy chain isotype and light chain type (classification)

We predicted the types of chains given heavy or light chains. Classes with fewer than 50 sequences were excluded. The target classes included: Heavy chain: IGHM, IGHD, IGHG, IGHA; Light chain type: IGK, IGL

#### Somatic hypermutation (SHM) frequency (regression)

We predicted the frequency of mutated nucleotides from the germline sequence in the junction region for heavy and light chains.

#### Junction length (regression)

We predicted the junction region length for the heavy and light chains. The junction region is the CDR3 plus the two flanking conserved amino acids.

#### Spike protein binding prediction (classification)

We predicted the binding or non-binding label of BCR heavy and light chain pair for the SARS-CoV-2 spike protein.

### Training and evaluation

To evaluate the performance of the embeddings on the prediction tasks, we trained the following simple machine-learning models.

### Classification tasks

We used the sklearn (Buitinck *et al*., 2013) support vector machine classifier (SVC) with RBF kernel and applied nested cross-validation to split the data into training, validation, and test sets with non-overlapping donors or studies and preserved class percentage (sklearn.model_selection.StratifiedGroupKFold). Five outer loops and four inner loops were used for gene usage and chain-type tasks, while four outer loops and three inner loops were used for specificity prediction due to a smaller dataset. We performed grid searches over the regularization parameter *C* of SVC ranging from 0.01 to 100 and selected the optimum based on the weighted-average F1 score based on the validation set performance. We finally evaluated the test set performance using the weighted-average F1 score, Matthews’ correlation coefficient, and balanced accuracy.

### Regression tasks

We chose the sklearn linear model with Lasso regularization (linear_model.Lasso) and used nested cross-validation (five outer loops and four inner loops) and split the dataset into training, validation, and test sets with non-overlapping samples. We performed grid searches over the regularization parameter α of the regressor ranging from 1e-6 to 1e-3 and selected the optimum based on the root mean square error (RMSE) score based on the validation set performance. We finally evaluated the test set performance by RMSE and adjusted R-square.

### Random baseline

To establish a baseline for the prediction performance, we randomly shuffled the labels to all data and repeated the same procedure described above for each task.

## RESULTS

### Evaluating BCR V(D)J amino acid sequence embeddings for sequence property and receptor specificity prediction tasks

We profiled the prediction performance of BCR embeddings on two types of tasks (Figure 1A). The first type of tasks consisted of fundamental BCR sequence properties, including V and J gene usage, chain type, somatic hypermutation frequency, and junction length. The second type of task focused on the receptor specificity to antigen. We used the specificity to SARS-CoV-2 spike protein as an example because of the data availability.

**Figure 1.**
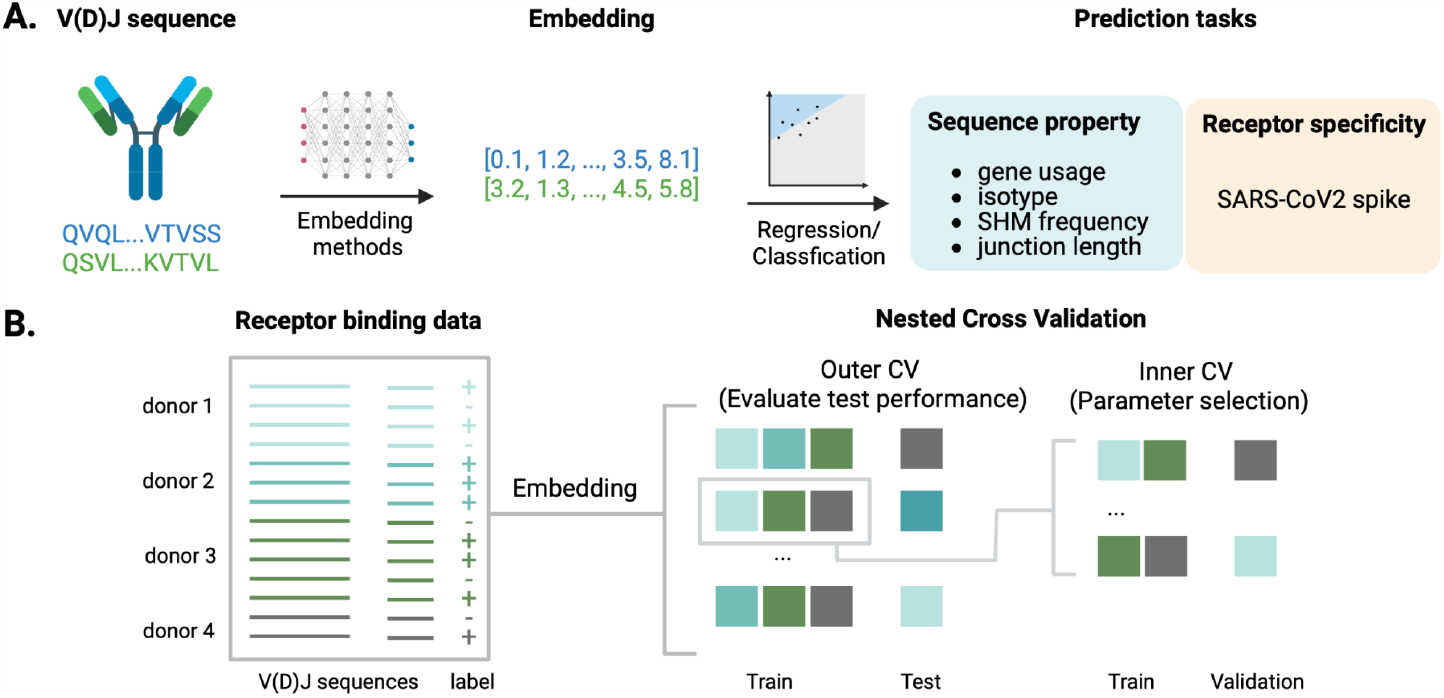
Benchmarking the performance of BCR amino acid embeddings on sequence property and receptor specificity prediction tasks. (A) BCR amino acid sequences were encoded by multiple embedding models and used to train supervised machine learning models for sequence properties (separately evaluated for heavy or light chains) or receptor specificity prediction. (B) Nested cross-validation (CV) evaluation of the embedding performance (using receptor specificity prediction tasks as an example). The inner CV loop was used to select optimal prediction model parameters, while the outer CV loop was used to evaluate the test performance. Each training, validation, and test split contained non-overlapping donors or studies.

To train and evaluate embedding models on these prediction tasks, we collected paired BCR heavy and light chain V(D)J sequence data from multiple data sources (Table 1). In total, the data consists of 0.87 million full-length heavy chains and 0.55 million full-length light chains. We also obtain unique CDR3 sequences from the data, which includes 0.79 million heavy chain CDR3, and 0.23 million light chain CDR3.

We chose five models to embed the BCR sequences: an amino acid 3-mer word2vec model (immune2vec), two pre-trained general protein language models (ESM2 and ProtT5), and two widely used encodings (physicochemical-based and amino acid frequency). We trained separate immune2vec models for heavy and light chains, as well as for full-length V(D)J and CDR3 sequences. For immune2vec, we used the suggested dimension of 100 from (Ostrovsky-Berman *et al*., 2021). In addition, to explore the effect of latent space dimensions on prediction performance, we trained immune2vec with dimensions ranging from 25 -1,000. Finally, we applied each model to embed both the full-length and CDR3 sequences of BCR heavy and light chains.

### Embeddings capture information on BCR sequence properties

To evaluate the ability of each of the embeddings to predict BCR sequence properties, we used the embeddings as input (Figure 2A-C, Figure 3A-B, Figure S1) to train supervised models on six classification tasks (V, J gene prediction for heavy and light chain, isotype, and light chain type) and four regression tasks (SHM frequency and junction length for heavy and light chain). Specifically, we used support vector machine classifiers for classification tasks and linear models with Lasso regularization for regression tasks. We chose simple prediction models over more expressive models to test whether the embeddings produced good high-level representations that relate to the underlying factors through simple dependencies (Bengio *et al*., 2014). This helps establish how useful the embeddings are for encoding more complicated, task-specific features that may rely heavily on basic properties.

**Figure 2.**
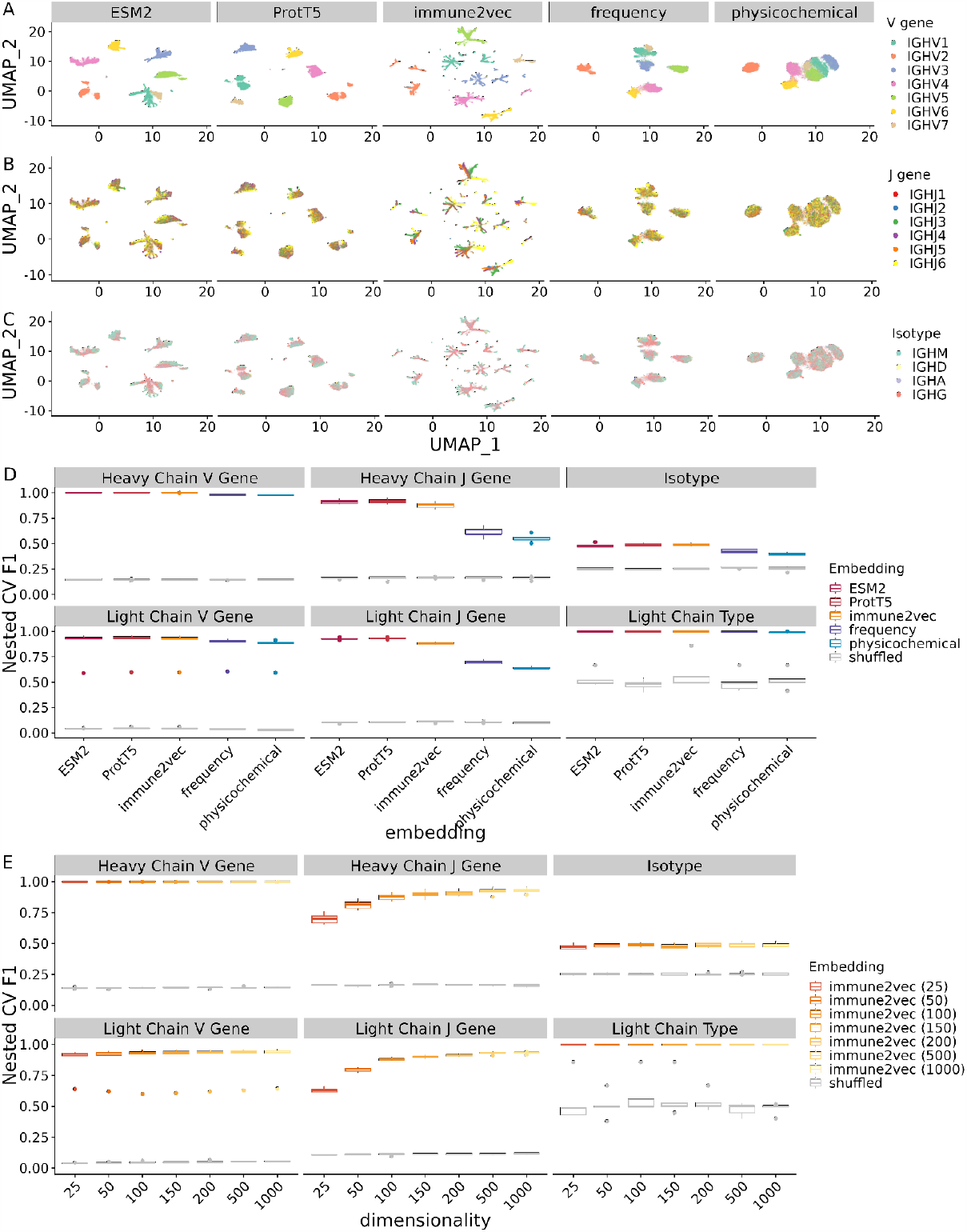
Performance of supervised models for sequence property classification tasks using BCR embeddings. (A-C) UMAP visualization of the BCR heavy chain embeddings, colored by V gene, J gene, and isotype, respectively. (D) Boxplot of prediction performance evaluated by the five outer folds of the nested cross-validation on sequence property classification tasks. The x- and y-axis show the type of embeddings under evaluation and the nested cross-validation weighted F1 score between the prediction and labels, respectively. Note that immune2vec models here have dimensions of 100 as recommended by the original paper, and separate immune2vec models were trained for the heavy and light sequences. The gray box plots indicate the performance of shuffled embeddings. (E) Effect of latent dimension size of immune2vec models on sequence property classification tasks.

**Figure 3.**
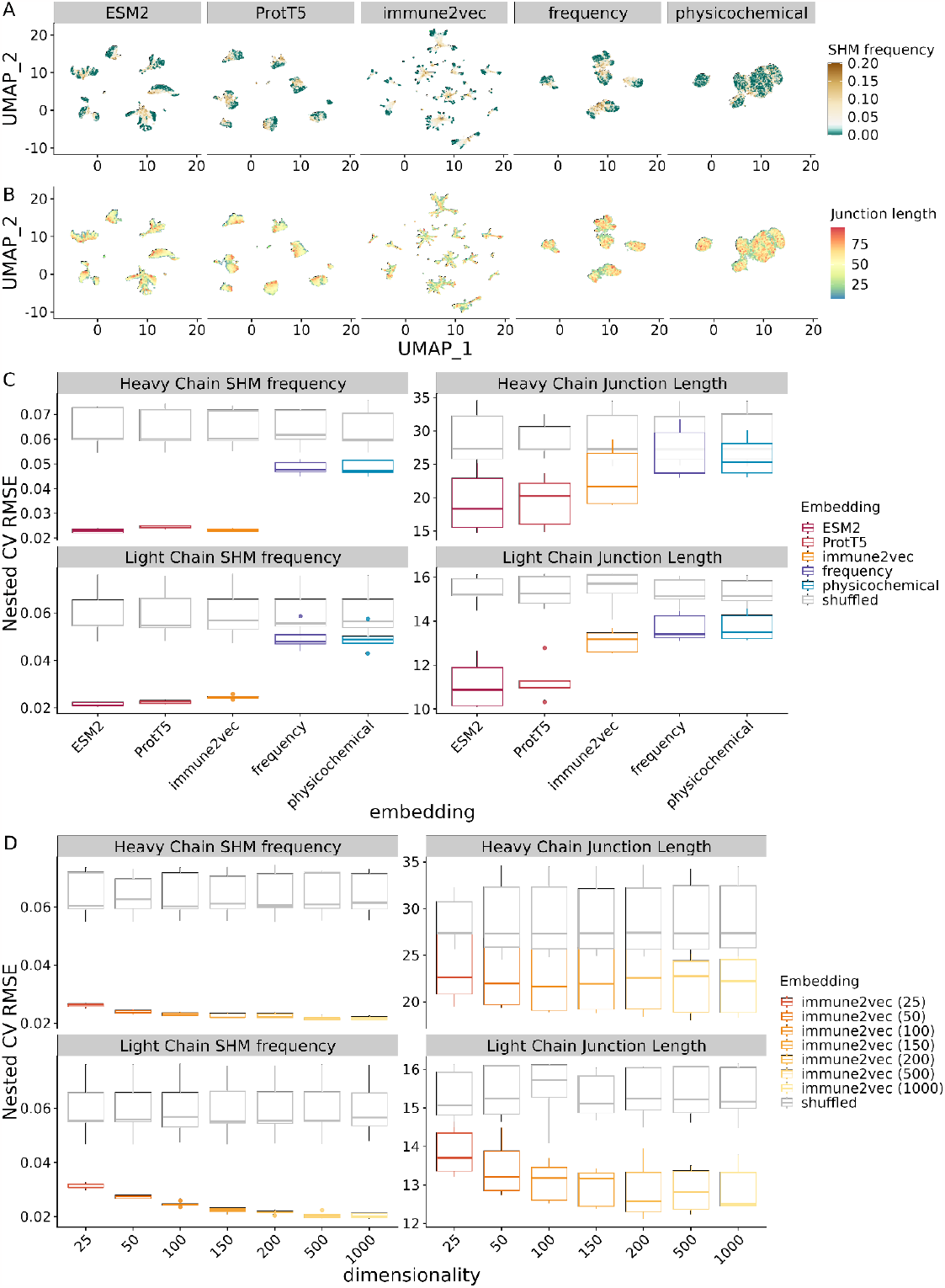
Performance of supervised models for sequence property regression tasks using BCR embeddings. (A-B) UMAP visualization of the BCR heavy chain embeddings, colored by somatic hypermutation frequency and junction length, respectively. (C) Boxplot of prediction performance evaluated by the five outer folds of the nested cross-validation on sequence property regression tasks. The x- and y-axis show the type of embeddings under evaluation and the nested cross-validation root mean square error between the prediction and labels, respectively. Note that immune2vec models here have dimensions of 100 as recommended by the original paper, and separate immune2vec models were trained for the heavy and light sequences. The gray box plots indicate the performance of shuffled embeddings. (D) Effect of latent dimension size of immune2vec models on sequence property regression tasks.

As is good practice in model selection (Pavlović *et al*., 2021), we performed nested cross-validation on the dataset to evaluate the supervised prediction model, with inner and outer loops for parameter selection and test set evaluation (Figure 1B). The train/validation/test splits were created with non-overlapping donors or studies. We also established a random baseline performance by repeating the nested cross-validation with randomly shuffled labels.

### Classification tasks

We evaluated the model performance using three balance-corrected accuracy measures: weighted F1 score, Matthew’s correlation coefficient (MCC), and balanced accuracy score (Figure 2D, Table S1). We chose these measures because of the class imbalance in the dataset; for example, the IGHV3 gene family has 100 times more sequences than the IGHV7 gene family. We found all embeddings performed significantly better than the randomly shuffled embeddings across all tasks. In addition, protein language models (ESM2, ProtT5, and immune2vec) outperformed baseline physicochemical and amino acid frequency embeddings by 3 -65 %, depending on the type of tasks. To further analyze the advantages of language models, we split the prediction tasks into three categories based on prediction performance: easy, moderate, and hard. All models performed well for easy tasks, including heavy and light chain V gene prediction, and light chain type prediction, suggesting saturation of modeling performance on these tasks. In contrast, all models performed poorly on the isotype prediction task. This is a hard task, likely due to the absence of the constant region sequence from input, so the information has to be inferred from the provided context. Most of the variation in embedding performance lies in moderate tasks: heavy and light chain J gene prediction tasks. ESM2, ProtT5, and immune2vec significantly outperformed the baseline physicochemical and frequency embeddings on the moderate tasks (Table S1). For example, the average nested CV weighted F1 scores for heavy chain J gene prediction tasks are 0.92, 0.92, and 0.88 for ESM2, ProtT5, and immune2vec, and 0.55, 0.61, 0.16 for physicochemical, frequency, and shuffled embedding, respectively.

To examine the effect of immune2vec dimensionality on prediction task accuracy, we compared the performance of the immune2vec with dimensions ranging from 25 to 1,000 (Figure 2E, Table S2). We found that the prediction performance increases as the embedding size increases for all tasks, especially for J gene prediction tasks. For example, the nested CV average F1 prediction performance for heavy chain J gene prediction steadily increases from 0.70 to 0.93 as immune2vec dimensions increase from 25 to 1,000. This indicates an increased capacity of the model to encode sequence information as the dimension increases.

### Regression Tasks

For the sequence property regression tasks, we measured the prediction performance using root mean square error (RMSE), adjusted R2 score (R2), and mean absolute error (MAE) (Figure 3C, Table S3). As with the sequence property classification tasks, we found that across all tasks, ESM2, ProtT5, and immune2vec performed better than the baseline physicochemical and frequency embeddings. Specifically, in the heavy chain somatic hypermutation (SHM) frequency prediction task, both language models and immune2vec reach around 0.02 in average nested CV RMSE, whereas the baseline embeddings had a higher RMSE of 0.05. We also noticed that among language models, ESM2 and ProtT5 performed better than immune2vec in predicting the junction lengths.

We also explored the effect of dimensionality on immune2vec prediction performance on regression tasks. Similar to the observations from classification tasks, immune2vec regression task prediction performance improved as the embedding size increased (Figure 3D, Table S4).

In summary, all five embeddings encode some level of sequence property information and perform much better than randomly shuffled embeddings. Immune2vec and protein language models capture more sequence property information than baseline frequency and physicochemical encodings. Larger immune2vec models can learn more information from the sequences and achieve better performance in sequence property prediction.

### Embeddings capture information to predict specificity to SARS-CoV-2 spike protein

We next evaluated how well BCR embeddings predict receptor specificity to the SARS-CoV-2 spike protein. To collect the datasets, we queried the Coronavirus Antibody Database (CoV-AbDab) for BCR sequences with binding information to SARS-CoV-2 wild-type spike protein. Because the dataset contains fewer non-binders, we sampled 1,000 random sequences from each donor of (Wang *et al*., 2023) dataset collected before the COVID-19 pandemic as non-binders, assuming that it is rare to find spike protein binders in pre-pandemic populations. In total, we obtained 15,538 sequences (55.7 % binders) from 34 donors/studies to evaluate the ability of embeddings to predict specificity to coronavirus spike protein.

Previous specificity prediction studies for BCR often focused on the CDR3 region of the heavy chain due to the limited availability of paired full-length V(D)J sequence data. Since the advent of single-cell technology, we have gained access to more paired full-length V(D)J sequences, making it possible to examine whether adding the region outside CDR3 and including paired light chain information helps with specificity prediction. We tested four different sequence inputs to the embedding methods, including paired full-length sequences (HL_Full), full-length heavy chain sequences (H_Full), paired CDR3 sequences (HL_CDR3), heavy chain CDR3 sequences (H_CDR3) (Figure 4A).

**Figure 4.**
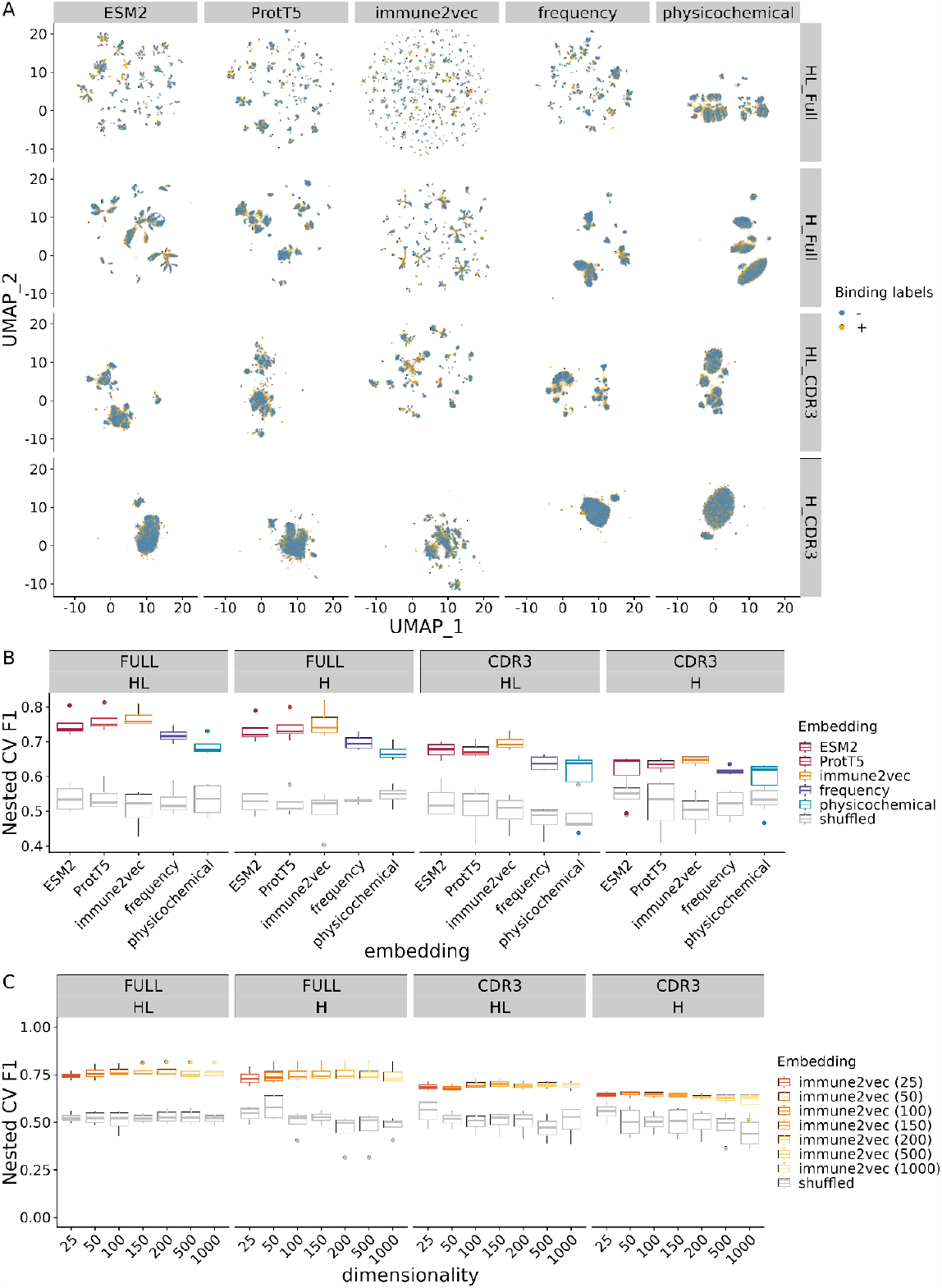
Performance of supervised models for receptor specificity tasks using BCR embeddings. (A) UMAP visualization of the BCR embeddings on various sequence inputs (HL_Full: full-length heavy and light chain, H_Full: full-length heavy chain, HL_CDR3: CDR3 heavy and light chain, H_CDR3: CDR3 heavy chain), colored by binding status (+: binders, -: non-binders). We trained separate immune2vec models for each sequence input type to embed the sequences. (B) Boxplot of prediction performance evaluated by the five outer folds of the nested cross-validation on receptor specificity tasks. The x- and y-axis show the embeddings under evaluation and the weighted F1 score between the prediction and labels, respectively. Note that immune2vec models here have dimensions of 100 as recommended by the original paper, and separate immune2vec models were trained for the heavy and light sequences as well as different BCR sequence inputs. The gray box plots indicate the performance of shuffled embeddings. (C) Effect of latent dimension size of immune2vec models on receptor specificity prediction task.

To evaluate the performance of the embeddings on different inputs, we trained the embedding models with corresponding input and predicted the binary binding labels using a simple support vector machine classifier. We applied nested cross-validation on the embeddings to compute the weighted F1, MCC, and balanced accuracy score of binding prediction (Figure 4B, Table S4). We found that all embeddings perform significantly better than the randomly shuffled embeddings. In addition, prediction performance improved as we included full-length and paired sequences across embedding methods (prediction performance: HL_Full > H_Full > HL_CDR3 > H_CDR3, Figure S2). This indicates that BCR regions outside CDR3, as well as the light chains, provide additional information on the specificity.

Comparing the embeddings, we found that ESM2, ProtT5, and immune2vec outperformed the baseline physicochemical and frequency embeddings. Interestingly, among language models, we found that pre-trained protein language models ESM2, and ProtT5 predicted no better than the smaller word embedding immune2vec models trained directly on BCR sequences. This may be due to the distinctive nature of BCR sequences compared to other protein sequences and raises the question of whether protein language models trained directly on BCR data could improve BCR modeling performance.

We also examined the relationship between receptor specificity prediction performance and the dimensionality of the immune2vec models. We found that the specificity prediction performance is not sensitive to the dimensionality of immune2vec (Figure 4C, Table S6). The performance first improves and then drops slightly as the dimensionality increases. The performance drop at higher dimensionality is more obvious for shorter sequences that contain less information for the embeddings to encode, which could lead to overfitting of the prediction model to noise given the relatively small dataset.

## DISCUSSION

In this study, we evaluated the performance of BCR sequence embedding methods, including immune2vec, ESM2, ProtT5, physicochemical, and amino acid frequency encodings, in predicting sequence properties and specificity to SARS-CoV-2 spike protein. We tested whether the embeddings produced good high-level representations that relate to the underlying biological properties through simple dependencies using linear models. We found that all embeddings encoded some information on sequence properties and specificity. Moreover, protein language models outperformed the baseline physicochemical and frequency embeddings across tasks. We also found that for sequence properties prediction, immune2vec models with higher latent dimensions learned more information from the sequences and performed better; however, for specificity prediction, the performance increased initially but stopped improving after the latent dimension became too high.

We also assessed the effect of sequence input for receptor specificity prediction. We found that using the full-length sequence as input and incorporating the light chain sequence improved the specificity prediction performance for all embedding methods, compared with using only CDR3 and heavy chain sequences. We also noticed a higher variance in prediction performance across the nested CV folds when using the full-length sequence. This may be due to the averaging procedure for generating sequence-level embeddings. While this is a common way to encode variable length information as a fixed length embedding (Ostrovsky-Berman *et al*., 2021; Lin *et al*., 2022; Elnaggar *et al*., 2020), averaging across the sequences may miss information present in only sections of a sequence. For example, the BCR sequence can be divided into four framework regions (FRs), which are relatively conserved and provide structural support, and three CDRs, the main determinants of antigen specificity. Full-length sequence input could introduce a higher percentage of irrelevant sequences, such as FRs, and averaging would lead to a higher noise level for these sequences. A potential solution could be training additional models to learn optimal ways to aggregate the embedding across the sequences (Detlefsen *et al*., 2022). We also noticed that the general protein language model performs similarly or slightly worse than the BCR-specific immune2vec model. A future direction will be using BCR-specific language models (Leem *et al*., 2022; Ruffolo *et al*., 2021) or fine-tuning the general protein language model using BCR sequences.

Training time can be important for some embeddings. For example, training an immune2vec model of dimension 100 on 0.8 million heavy chain full-length sequences took about 2.5 hours on a single CPU core from a standard HPC node using less than 5G memory. For protein language models ESM2 and ProtT5, we downloaded the pre-trained models. The sequence embedding time was very different between immune2vec and the protein language model. The immune2vec model embeds 0.8 million heavy chain sequences in about 30 minutes on a single CPU core from a standard HPC node with 50G memory requested, while ESM2 and ProtT5 took 1 -2 days with the same computing resource setting.

Some limitations of our study are the small sample size for the receptor specificity task and that our data is aggregated over a limited set of studies and patients, which can miss some binding modes and introduce batch effects. To mitigate potential batch effects, we used the nested CV framework. The other main data-related limitation is the label imbalance. For example, CoV-AbDab contains mostly binders, and we added random sequences from the pre-pandemic samples as negatives. We used an even partition across each fold to ensure that this does not create a biased model, which reduces the sample size. We also used evaluation metrics on the classifiers that would account for this data limitation, as MCC and weighted-average F1 scores are known to be robust to label imbalance.

In summary, we found that language models outperformed traditional amino acid encodings. The BCR-specific immune2vec model slightly outperformed general protein language models in specificity prediction for SARS-CoV-2 spike protein, and incorporating full-length and light chain sequences improved the specificity prediction performance. The findings give insights into future studies using BCR embeddings for downstream prediction applications.

## DATA AVAILABILITY

All data were from public sources as listed in Table 1. The code is available on bitbucket (https://bitbucket.org/kleinstein/projects/src/master/Wang2023/).

## ACKNOWLEDGEMENTS

We thank Ronen Basri for the method discussion and Kenneth Hoehn for data processing.

