## Supplemental Figure S1 - S2, Table S1 - S6 for "Language model-based B cell receptor sequence embeddings can effectively encode receptor specificity"

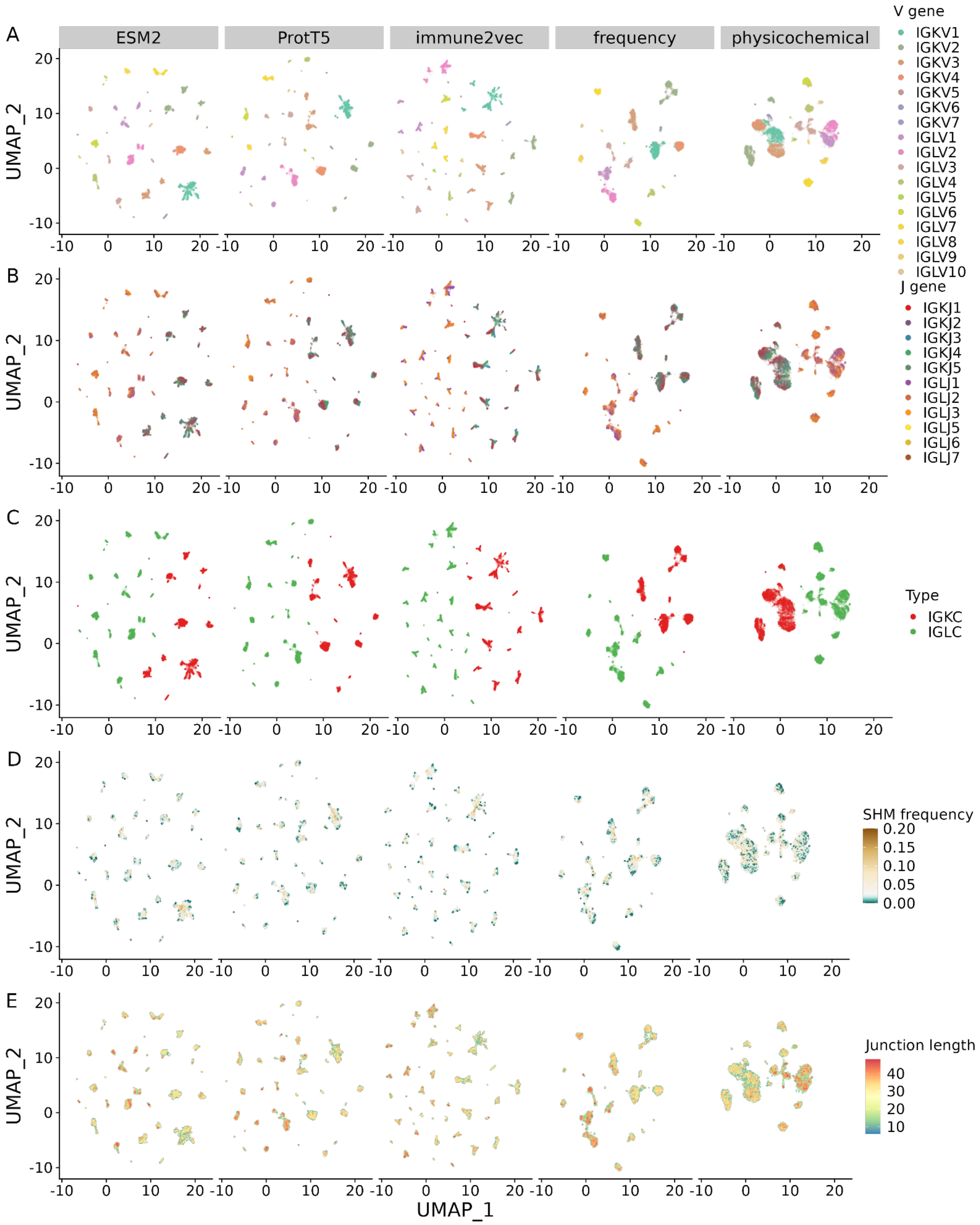


**Figure S1. UMAP visualization of BCR light chain embeddings colored by** (A) V gene family, (B) J gene family, (C) light chain type, (D) Somatic hypermutation frequency, (E) junction length.

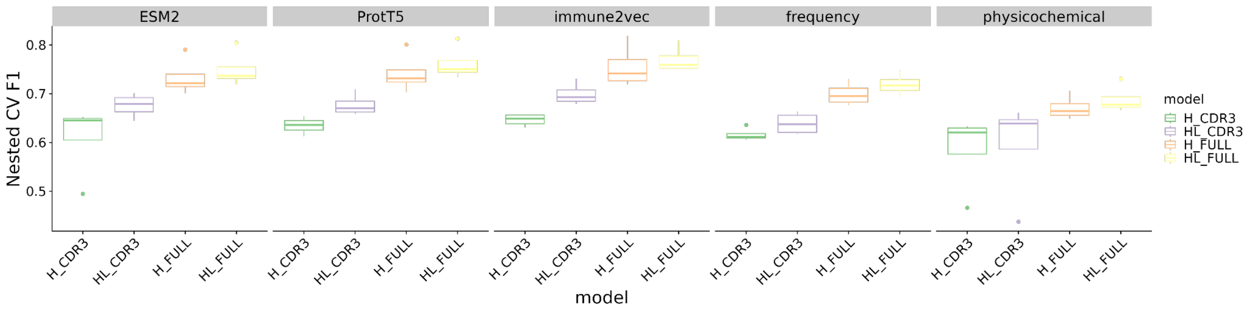


**Figure S2. Receptor specificity prediction performance across embeddings and sequence inputs**. H_CDR3: CDR3 heavy chain embedding, HL_CDR3: paired CDR3 heavy and light chain embedding concatenated, H_FULL: full-length heavy chain embedding, HL_FULL: paired full-length heavy and light chain embedding.

| Task | Embedding | F1 | MCC | ACC |
|---|---|---|---|---|
| Heavy Chain V Gene | ESM2 | 0.99976 | 0.99971 | 0.99976 |
| Heavy Chain V Gene | ProtT5 | 0.99987 | 0.99985 | 0.99987 |
| Heavy Chain V Gene | immune2vec | 0.99991 | 0.99990 | 0.99991 |
| Heavy Chain V Gene | frequency | 0.97772 | 0.97379 | 0.97771 |
| Heavy Chain V Gene | physicochemical | 0.97603 | 0.97181 | 0.97602 |
| Heavy Chain J Gene | ESM2 | 0.91652 | 0.89979 | 0.91633 |
| Heavy Chain J Gene | ProtT5 | 0.91850 | 0.90192 | 0.91835 |
| Heavy Chain J Gene | immune2vec | 0.87761 | 0.85220 | 0.87579 |
| Heavy Chain J Gene | frequency | 0.60902 | 0.53365 | 0.61030 |
| Heavy Chain J Gene | physicochemical | 0.55156 | 0.46846 | 0.55574 |
| Isotype | ESM2 | 0.48162 | 0.28877 | 0.48172 |
| Isotype | ProtT5 | 0.48803 | 0.29502 | 0.48818 |
| Isotype | immune2vec | 0.49020 | 0.30190 | 0.49170 |
| Isotype | frequency | 0.42697 | 0.21277 | 0.42684 |
| Isotype | physicochemical | 0.39883 | 0.17982 | 0.39836 |
| Light Chain V Gene | ESM2 | 0.86924 | 0.86787 | 0.86385 |
| Light Chain V Gene | ProtT5 | 0.87409 | 0.87211 | 0.86834 |
| Light Chain V Gene | immune2vec | 0.87079 | 0.86787 | 0.86469 |
| Light Chain V Gene | frequency | 0.84714 | 0.84016 | 0.83998 |
| Light Chain V Gene | physicochemical | 0.83482 | 0.82673 | 0.82793 |

| | | | | |
|---|---|---|---|---|
| Light Chain J Gene | ESM2 | 0.92711 | 0.91779 | 0.92741 |
| Light Chain J Gene | ProtT5 | 0.93189 | 0.92294 | 0.93201 |
| Light Chain J Gene | immune2vec | 0.88107 | 0.86609 | 0.88179 |
| Light Chain J Gene | frequency | 0.70072 | 0.66166 | 0.70093 |
| Light Chain J Gene | physicochemical | 0.63796 | 0.59045 | 0.63679 |
| Light Chain Type | ESM2 | 1.00000 | 0.80000 | 1.00000 |
| Light Chain Type | ProtT5 | 1.00000 | 0.80000 | 1.00000 |
| Light Chain Type | immune2vec | 1.00000 | 0.80000 | 1.00000 |
| Light Chain Type | frequency | 0.99691 | 0.79385 | 0.99692 |
| Light Chain Type | physicochemical | 0.99293 | 0.78587 | 0.99293 |

**Table S1. Performance of BCR embeddings on sequence property classification tasks.** Nest cross-validation was performed to evaluate the average weighted F1 score (F1), Matthew's correlation coefficient (MCC), and balanced accuracy (ACC) across the outer loops.

| Dimensionality | Task | F1 | MCC | ACC |
|---|---|---|---|---|
| 25 | Heavy Chain V Gene | 0.99964 | 0.99957 | 0.99964 |
| 50 | Heavy Chain V Gene | 0.99988 | 0.99986 | 0.99988 |
| 100 | Heavy Chain V Gene | 0.99991 | 0.99990 | 0.99991 |
| 150 | Heavy Chain V Gene | 0.99991 | 0.99990 | 0.99991 |
| 200 | Heavy Chain V Gene | 0.99992 | 0.99991 | 0.99992 |
| 500 | Heavy Chain V Gene | 0.99991 | 0.99989 | 0.99991 |
| 1000 | Heavy Chain V Gene | 0.99998 | 0.99998 | 0.99998 |
| 25 | Heavy Chain J Gene | 0.70238 | 0.64558 | 0.70426 |
| 50 | Heavy Chain J Gene | 0.81401 | 0.77564 | 0.81192 |
| 100 | Heavy Chain J Gene | 0.87761 | 0.85220 | 0.87579 |
| 150 | Heavy Chain J Gene | 0.89795 | 0.87720 | 0.89726 |
| 200 | Heavy Chain J Gene | 0.91067 | 0.89219 | 0.90998 |
| 500 | Heavy Chain J Gene | 0.92328 | 0.90763 | 0.92280 |
| 1000 | Heavy Chain J Gene | 0.92885 | 0.91427 | 0.92852 |
| 25 | Isotype | 0.47532 | 0.28452 | 0.47687 |
| 50 | Isotype | 0.48663 | 0.29829 | 0.48801 |

| | | | | |
|---|---|---|---|---|
| 100 | Isotype | 0.49020 | 0.30190 | 0.49170 |
| 150 | Isotype | 0.48122 | 0.29626 | 0.48429 |
| 200 | Isotype | 0.48703 | 0.29762 | 0.48912 |
| 500 | Isotype | 0.48670 | 0.29918 | 0.48737 |
| 1000 | Isotype | 0.48855 | 0.30091 | 0.48905 |
| 25 | Light Chain V Gene | 0.86612 | 0.85616 | 0.85717 |
| 50 | Light Chain V Gene | 0.86701 | 0.86078 | 0.85945 |
| 100 | Light Chain V Gene | 0.87079 | 0.86787 | 0.86469 |
| 150 | Light Chain V Gene | 0.87447 | 0.87092 | 0.86804 |
| 200 | Light Chain V Gene | 0.87713 | 0.87261 | 0.87021 |
| 500 | Light Chain V Gene | 0.88128 | 0.87614 | 0.87404 |
| 1000 | Light Chain V Gene | 0.88480 | 0.87821 | 0.87699 |
| 25 | Light Chain J Gene | 0.63304 | 0.59067 | 0.63793 |
| 50 | Light Chain J Gene | 0.79688 | 0.77137 | 0.79812 |
| 100 | Light Chain J Gene | 0.88107 | 0.86609 | 0.88179 |
| 150 | Light Chain J Gene | 0.90079 | 0.88799 | 0.90117 |
| 200 | Light Chain J Gene | 0.91214 | 0.90049 | 0.91222 |
| 500 | Light Chain J Gene | 0.92960 | 0.92019 | 0.92959 |
| 1000 | Light Chain J Gene | 0.93278 | 0.92403 | 0.93293 |
| 25 | Light Chain Type | 1.00000 | 0.80000 | 1.00000 |
| 50 | Light Chain Type | 1.00000 | 0.80000 | 1.00000 |
| 100 | Light Chain Type | 1.00000 | 0.80000 | 1.00000 |
| 150 | Light Chain Type | 1.00000 | 0.80000 | 1.00000 |
| 200 | Light Chain Type | 1.00000 | 0.80000 | 1.00000 |
| 500 | Light Chain Type | 1.00000 | 0.80000 | 1.00000 |
| 1000 | Light Chain Type | 1.00000 | 0.80000 | 1.00000 |

**Table S2. Changes in immune2vec embeddings performance with respect to dimensionality on sequence property classification tasks.** Immune2vec models were trained with different dimensions (25 - 1000) to evaluate the effect of dimensionality on prediction performance. Nested cross-validation was performed to evaluate the average weighted F1 score (F1), Matthew's correlation coefficient (MCC), and balanced accuracy (ACC) across the outer loops.

| Task | Embedding | RMSE | R2 | MAE |
|---|---|---|---|---|
| Heavy Chain SHM frequency | ESM2 | 0.02296 | 0.85567 | 0.01740 |
| Heavy Chain SHM frequency | ProtT5 | 0.02445 | 0.83653 | 0.01861 |
| Heavy Chain SHM frequency | immune2vec | 0.02313 | 0.85605 | 0.01765 |
| Heavy Chain SHM frequency | frequency | 0.04842 | 0.37989 | 0.03937 |
| Heavy Chain SHM frequency | physicochemical | 0.04838 | 0.38469 | 0.03919 |
| Heavy Chain Junction Length | ESM2 | 19.34430 | -6.23156 | 17.22838 |
| Heavy Chain Junction Length | ProtT5 | 19.38826 | -0.20392 | 17.05321 |
| Heavy Chain Junction Length | immune2vec | 23.02751 | -9.02568 | 20.04956 |
| Heavy Chain Junction Length | frequency | 26.35413 | -11.23459 | 23.34738 |
| Heavy Chain Junction Length | physicochemical | 26.07709 | -1.19957 | 22.96459 |
| Light Chain SHM frequency | ESM2 | 0.02155 | 0.85010 | 0.01630 |
| Light Chain SHM frequency | ProtT5 | 0.02260 | 0.83667 | 0.01725 |
| Light Chain SHM frequency | immune2vec | 0.02459 | 0.80612 | 0.01850 |
| Light Chain SHM frequency | frequency | 0.04976 | 0.25127 | 0.03907 |
| Light Chain SHM frequency | physicochemical | 0.04946 | 0.25512 | 0.03887 |
| Light Chain Junction Length | ESM2 | 11.13957 | -0.78333 | 9.69484 |
| Light Chain Junction Length | ProtT5 | 11.26954 | -0.75678 | 9.76896 |
| Light Chain Junction Length | immune2vec | 13.09683 | -1.39173 | 11.95008 |
| Light Chain Junction Length | frequency | 13.69778 | -1.58550 | 12.68878 |
| Light Chain Junction Length | physicochemical | 13.74507 | -1.75419 | 12.75006 |

**Table S3. Performance of BCR embeddings on sequence property regression tasks.** Nest cross-validation was performed to evaluate the average root mean square error (RMSE), adjusted R2 (R2), and mean absolute error (MAE) across the outer loops.

| Dimensionality | Task | RMSE | R2 | MAE |
|---|---|---|---|---|
| 25 | Heavy Chain SHM frequency | 0.02620 | 0.81669 | 0.02016 |
| 50 | Heavy Chain SHM frequency | 0.02387 | 0.84633 | 0.01815 |
| 100 | Heavy Chain SHM frequency | 0.02313 | 0.85605 | 0.01765 |
| 150 | Heavy Chain SHM frequency | 0.02264 | 0.86081 | 0.01726 |
| 200 | Heavy Chain SHM frequency | 0.02263 | 0.86278 | 0.01722 |

| | | | | |
|---|---|---|---|---|
| 500 | Heavy Chain SHM frequency | 0.02177 | 0.87329 | 0.01644 |
| 1000 | Heavy Chain SHM frequency | 0.02182 | 0.87219 | 0.01645 |
| 25 | Heavy Chain Junction Length | 24.10200 | -9.78745 | 21.13150 |
| 50 | Heavy Chain Junction Length | 23.39717 | -9.14411 | 20.36449 |
| 100 | Heavy Chain Junction Length | 23.02751 | -9.02568 | 20.04956 |
| 150 | Heavy Chain Junction Length | 22.85182 | -7.77051 | 19.75448 |
| 200 | Heavy Chain Junction Length | 22.85676 | -8.05100 | 19.81380 |
| 500 | Heavy Chain Junction Length | 21.97517 | -0.54174 | 18.83146 |
| 1000 | Heavy Chain Junction Length | 22.38640 | -7.71062 | 19.23070 |
| 25 | Light Chain SHM frequency | 0.03126 | 0.69164 | 0.02395 |
| 50 | Light Chain SHM frequency | 0.02735 | 0.75988 | 0.02061 |
| 100 | Light Chain SHM frequency | 0.02459 | 0.80612 | 0.01850 |
| 150 | Light Chain SHM frequency | 0.02236 | 0.83817 | 0.01683 |
| 200 | Light Chain SHM frequency | 0.02158 | 0.85161 | 0.01624 |
| 500 | Light Chain SHM frequency | 0.02049 | 0.86538 | 0.01534 |
| 1000 | Light Chain SHM frequency | 0.02029 | 0.86732 | 0.01514 |
| 25 | Light Chain Junction Length | 13.85732 | -1.85679 | 12.91240 |
| 50 | Light Chain Junction Length | 13.43479 | -1.61517 | 12.38213 |
| 100 | Light Chain Junction Length | 13.09683 | -1.39173 | 11.95008 |
| 150 | Light Chain Junction Length | 12.94980 | -1.33768 | 11.76980 |
| 200 | Light Chain Junction Length | 12.85761 | -1.25761 | 11.65768 |
| 500 | Light Chain Junction Length | 12.85821 | -1.35267 | 11.60367 |
| 1000 | Light Chain Junction Length | 12.90942 | -1.52857 | 11.83083 |

**Table S4. Changes in immune2vec embeddings performance with respect to dimensionality on sequence property regression tasks.** Immune2vec models were trained with different dimensions (25 - 1000) to evaluate the effect of dimensionality on prediction performance. Nested cross-validation was performed to evaluate the average root mean square error (RMSE), adjusted R2 (R2), and mean absolute error (MAE) across the outer loops.

| Embedding | Model | Sequence | F1 | MCC | ACC | AUROC |
|---|---|---|---|---|---|---|
| ESM2 | H | CDR3 | 0.60926 | 0.17603 | 0.59803 | 0.59803 |
| ProtT5 | H | CDR3 | 0.63451 | 0.19168 | 0.60829 | 0.60829 |

| | | | | | | |
|---|---|---|---|---|---|---|
| immune2vec | H | CDR3 | 0.64611 | 0.19681 | 0.61062 | 0.61062 |
| frequency | H | CDR3 | 0.61605 | 0.12876 | 0.57148 | 0.57148 |
| physicochemical | H | CDR3 | 0.58512 | 0.12100 | 0.56746 | 0.56746 |
| ESM2 | H | FULL | 0.73344 | 0.34726 | 0.68567 | 0.68567 |
| ProtT5 | H | FULL | 0.74161 | 0.36610 | 0.69588 | 0.69588 |
| immune2vec | H | FULL | 0.75540 | 0.38084 | 0.70213 | 0.70213 |
| frequency | H | FULL | 0.69916 | 0.28339 | 0.65464 | 0.65464 |
| physicochemical | H | FULL | 0.67093 | 0.23082 | 0.62642 | 0.62642 |
| ESM2 | HL | CDR3 | 0.67587 | 0.26867 | 0.65034 | 0.65034 |
| ProtT5 | HL | CDR3 | 0.67696 | 0.27244 | 0.65228 | 0.65228 |
| immune2vec | HL | CDR3 | 0.69919 | 0.29799 | 0.66538 | 0.66538 |
| frequency | HL | CDR3 | 0.63892 | 0.20820 | 0.61718 | 0.61718 |
| physicochemical | HL | CDR3 | 0.59417 | 0.16532 | 0.59236 | 0.59236 |
| ESM2 | HL | FULL | 0.74949 | 0.38407 | 0.70602 | 0.70602 |
| ProtT5 | HL | FULL | 0.76205 | 0.41351 | 0.72184 | 0.72184 |
| immune2vec | HL | FULL | 0.77024 | 0.41612 | 0.72064 | 0.72064 |
| frequency | HL | FULL | 0.71882 | 0.32801 | 0.67879 | 0.67879 |
| physicochemical | HL | FULL | 0.68823 | 0.26771 | 0.64583 | 0.64583 |

**Table S5. Performance of BCR embeddings on receptor specificity prediction tasks.** Nest cross-validation was performed to evaluate the average weighted F1 score (F1), Matthew's correlation coefficient (MCC), balanced accuracy (ACC), and area under the receiver operating characteristics (AUROC) across the outer loops.

| Dimensionality | Model | Sequence | F1 | MCC | ACC | AUROC |
|---|---|---|---|---|---|---|
| 25 | H | CDR3 | 0.64503 | 0.17744 | 0.59878 | 0.59878 |
| 50 | H | CDR3 | 0.64911 | 0.19428 | 0.60910 | 0.60910 |
| 100 | H | CDR3 | 0.64611 | 0.19681 | 0.61062 | 0.61062 |
| 150 | H | CDR3 | 0.64305 | 0.19791 | 0.61146 | 0.61146 |
| 200 | H | CDR3 | 0.63477 | 0.18228 | 0.60228 | 0.60228 |
| 500 | H | CDR3 | 0.62783 | 0.17731 | 0.59921 | 0.59921 |
| 1000 | H | CDR3 | 0.60506 | 0.15641 | 0.58631 | 0.58631 |

| | | | | | | |
|---|---|---|---|---|---|---|
| 25 | HL | CDR3 | 0.68858 | 0.27881 | 0.65513 | 0.65513 |
| 50 | HL | CDR3 | 0.68074 | 0.27538 | 0.65382 | 0.65382 |
| 100 | HL | CDR3 | 0.69919 | 0.29799 | 0.66538 | 0.66538 |
| 150 | HL | CDR3 | 0.70371 | 0.30946 | 0.67176 | 0.67176 |
| 200 | HL | CDR3 | 0.69318 | 0.29069 | 0.66194 | 0.66194 |
| 500 | HL | CDR3 | 0.69650 | 0.30191 | 0.66825 | 0.66825 |
| 1000 | HL | CDR3 | 0.69282 | 0.29736 | 0.66579 | 0.66579 |
| 25 | H | FULL | 0.73577 | 0.34113 | 0.68307 | 0.68307 |
| 50 | H | FULL | 0.74920 | 0.36449 | 0.69366 | 0.69366 |
| 100 | H | FULL | 0.75540 | 0.38084 | 0.70213 | 0.70213 |
| 150 | H | FULL | 0.75912 | 0.38889 | 0.70594 | 0.70594 |
| 200 | H | FULL | 0.75994 | 0.39183 | 0.70800 | 0.70800 |
| 500 | H | FULL | 0.75667 | 0.39011 | 0.70779 | 0.70779 |
| 1000 | H | FULL | 0.74906 | 0.37547 | 0.70085 | 0.70085 |
| 25 | HL | FULL | 0.74485 | 0.36293 | 0.69420 | 0.69420 |
| 50 | HL | FULL | 0.76020 | 0.39052 | 0.70657 | 0.70657 |
| 100 | HL | FULL | 0.77024 | 0.41612 | 0.72064 | 0.72064 |
| 150 | HL | FULL | 0.76938 | 0.41248 | 0.71807 | 0.71807 |
| 200 | HL | FULL | 0.77004 | 0.41893 | 0.72268 | 0.72268 |
| 500 | HL | FULL | 0.76096 | 0.40433 | 0.71623 | 0.71623 |
| 1000 | HL | FULL | 0.76128 | 0.40904 | 0.71960 | 0.71960 |

**Table S6. Changes in immune2vec embeddings performance with respect to dimensionality on receptor specificity prediction tasks.** Immune2vec models were trained with different dimensions (25 - 1000) and different sequence inputs to evaluate the effect of dimensionality on prediction performance. Nested cross-validation was performed to evaluate the average weighted F1 score (F1), Matthew's correlation coefficient (MCC), balanced accuracy (ACC), and area under the receiver operating characteristics (AUROC) across the outer loops.
